# Genetic and anatomical determinants of olfaction in dogs and wild canids

**DOI:** 10.1101/2024.04.15.589487

**Authors:** Alice Mouton, Deborah Bird, Gang Li, Brent A. Craven, Jonathan M. Levine, Marco Morselli, Matteo Pellegrini, Blaire Van Valkenburgh, Robert K. Wayne, William J. Murphy

## Abstract

Understanding the anatomical and genetic basis of complex phenotypic traits has long been a challenge for biological research. Domestic dogs offer a compelling model as they demonstrate more phenotypic variation than any other vertebrate species. Dogs have been intensely selected for specific traits and abilities, directly or indirectly, over the past 15,000 years since their initial domestication from the gray wolf. Because olfaction plays a central role in critical tasks, such as the detection of drugs, diseases, and explosives, as well as human rescue, we compared relative olfactory capacity across dog breeds and assessed changes to the canine olfactory system resulting from domestication. We conducted a cross-disciplinary survey of olfactory anatomy, olfactory receptor (OR) gene variation, and OR gene expression in domestic dogs. Through comparisons to their closest wild canid relatives, the gray wolf and coyote, we show that domestic dogs might have lost functional OR genes commensurate with a documented reduction in nasal morphology during domestication. Critically, within domestic dogs alone, we found no genetic or morphological profile shared among functional or genealogical breed groupings, such as scent hounds, that might indicate evidence of any human-directed selection for enhanced olfaction. Instead, our results suggest that superior scent detection dogs likely owe their success to advantageous behavioral traits and training rather than an “olfactory edge” provided by morphology or genes.

## Introduction

The olfactory acumen of the domestic dog (*Canis lupus familiaris*), particularly in breeds such as the bloodhound, is well established in popular lore and legend (Pemberton 2013; Worboys et al. 2018) (1, 2). In practice, dogs perform critical scent detection tasks, tracking missing persons, identifying individuals with diseases, (e.g., cancer, COVID-19) and locating explosives as well as cryptic and endangered species in the field (Helton 2009; Rooney et al. 2013; Beebe et al. 2016; Wackermannová et al. 2016; Gottwald et al. 2020; Jendrny et al. 2020; Dargan and Forbes 2021; Grimm-Seyfarth et al. 2021; Jendrny et al. 2021). Canine olfactory detection has even been used as admissible evidence in a court of law (Smith 2021). Aside from genuine feats and murkier legend, comparative studies on mammalian olfactory systems have ranked the dog’s olfactory capacity high relative to most other sampled species. This is the case when comparing olfactory anatomy as well as olfactory receptor (OR) gene repertoire size, a metric that has been correlated with scent discrimination performance ((Laska and Shepherd 2007; Rizvanovic et al. 2013). Behavioral tests of odorant discrimination and detection thresholds further reinforce the domestic dog’s standing as a superior smeller among mammals (Lauruschkus 1942; Marshall et al. 1981; Pihlström et al. 2005; Walker et al. 2006; Niimura et al. 2015; Bird et al. 2018), even if studies comparing detection performance across dog breeds arrive at no clear consensus on exceptional breeds (Jezierski et al. 2014; Hall et al. 2015; Polgár et al. 2016). Because an evolutionary perspective is absent from these studies, two important aspects of dog olfactory systems remain unclear.

First, the role artificial selection has played in shaping the dog’s olfactory system during its domestication from its wolf ancestor has yet to be examined. Second, the general assumption that concerted artificial selection has resulted in meaningful olfactory differences between dog breeds and breed groupings (e.g., scent breeds) has not been tested. Beginning more than 15,000 years ago, humans began domesticating individual wolves (*Canis lupus*) as commensal or companion animals, a process that would eventually shift to selective breeding for type (Bergström et al. 2020; Morrill et al. 2022). This process has resulted in a new canid lineage with a stunning breadth of variation in form and behavior, exceeding that of their common ancestor, the wolf. Here we investigate the influence of domestication and breed formation on the olfactory morphology, OR gene repertoires, and gene expression of the domestic dog.

Previous studies have established that mammals rely to varying degrees on olfaction for survival, as evidenced in losses and gains to species’ olfactory systems over time (Niimura and Nei 2007; Hayden et al. 2010; Bird et al. 2018). The OR gene superfamily, the largest gene family in terrestrial mammals (Buck and Axel 1991; Hayden et al. 2010), is variably represented in each species’ genomes (e.g., humans have a subgenome of 396 functional OR genes, whereas African elephants have 1,948). In parallel with OR gene repertoires, olfactory nasal morphology varies markedly across species and reflects selective sensory pressures (Van Valkenburgh et al. 2011; Bird et al. 2018; Bird et al. 2020).

In this light, we predict that the domestic dog, over time, by trading predatory behavior for reliance on human-provided food (Cannon et al. 1999; Arendt et al. 2016; Vonholdt and Driscoll 2017), underwent degeneration of olfactory function relative to its closest living canid relative, the gray wolf. On the other hand, if we accept the widespread assumption that certain dogs, such as scent hounds, have enhanced olfactory traits due to directed artificial selection, we might expect that these dogs have recovered some of this loss through copy number expansion (Gazit and Terkel 2003; Quignon et al. 2012; Pemberton 2013; Greenberg 2017). Moreover, we predict that ancient dogs, a grouping comprised of the most genetically divergent breeds (e.g. dingo, basenji, Siberian husky, Afghan, saluki) that show strong evidence of admixture with wolves after domestication, may retain wolf-like attributes including an enhanced olfactory system (Vonholdt et al. 2010; Freedman et al. 2014; Parker et al. 2017). A recent comparative morphological study of the cribriform plate (CP), the quantifiable bony imprint of olfactory nerves entering the brain from the nose, revealed that domestic dogs had a reduced olfactory skeleton relative to the gray wolf and that, contrary to claims of breeders, scent breeds show no enhanced olfactory phenotypes relative to other breeds (Bird et al. 2021)) (Fig. 1A; SI Appendix, Movie S1,2). No comprehensive molecular genetic analysis has been performed comparing OR gene repertoires across dog breeds and closely related wild canids. We hypothesized that OR gene repertoire size and gene expression levels might be more sensitive than morphology alone to directional selection for enhanced scent detection. Here we report the results of our cross-disciplinary study that combined morphology and molecular genetics from 56 domestic dog breeds, the gray wolf, and the coyote to better understand the evolutionary dynamics of olfactory systems in domestic dogs and their closest wild canid relatives.

**Figure 1.**
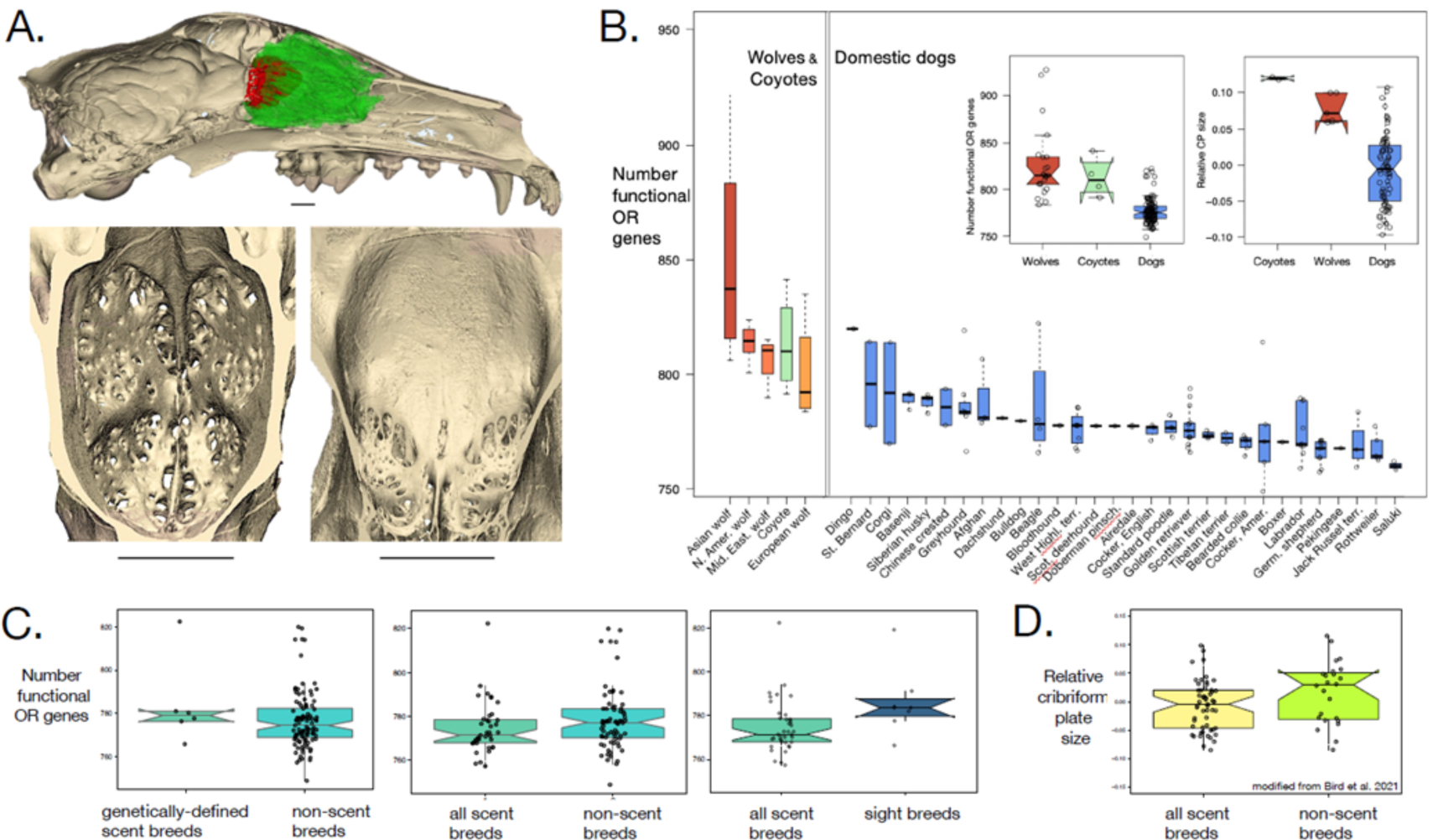
Comparative olfactory morphology and genomics in wild canids and dog breeds. A. Morphological metric, cribriform plate (CP) in skull matrix; Top, Borzoi half skull, sagittal view; red, CP; green, olfactory turbinal bones. Bottom, CP, posterior view, showing the relative presence of foramina (holes) for olfactory nerve passage in the gray wolf (left) and Pekingese (right). Scale bars, 10mm. B. Number of functional olfactory receptor genes (FORG) in wolves (orange), coyotes (green), and domestic dog breeds (blue) in descending order. Left inset, Domestic dogs have a smaller FORG repertoire than gray wolves and combined wolves and coyotes (p<0.001). Right inset, relative CP (RelCP) size in dogs is, on average, smaller than in wild canids (p<0.001). C. Left, FORG count in genetically-defined scent breeds is not significantly different from non-scent breeds (p=0.41). Middle, Mean FORG count for all scent breeds is not significantly different from that of non-scent breeds (p=0.16) and (right) sight breeds (p=0.08). D, RelCP size is no different between scent and non-scent breeds (n=46, p=0.12). Box plots: midline is median, whiskers are 5%-95% percentile.

## Results

### Olfactory genomics

To test whether losses in olfactory subgenomes accompanied the transition from wolf to dog, we quantified functional olfactory receptor gene (FORG) copy numbers for individual dogs, breeds, and breed groupings (e.g., ancient vs. modern, scent vs. non-scent, American Kennel Club: AKC, etc.) and for gray wolf populations and coyotes (*Canis latrans*) (SI Appendix, Dataset S1, S2). Overall, FORG repertoire size varies considerably across our sample of domestic dogs and their wild canid relatives (ANOVA, p < 0.001). When comparing our domestic dog and wild canid samples, between-group variation is larger than the within-group variation, and dogs, on average, have significantly fewer FORG than either the wolves alone or the wolves and coyotes combined (t-test, p < 0.001) (Fig.1B; SI Appendix, Table S1).

Within the entire wild canid sample, the number of FORG does not differ significantly between the gray wolf and coyote (n = 23, 4, t-test: p = 0.4). Within our gray wolf sample alone, the number of FORG ranges from 784 to 927, and the variance between wolf populations is significant (ANOVA, p = 0.025) (Fig. 1B; SI AppendixTable S1, Dataset S1, S2). Asian (Chinese and Mongolian) wolves (n = 9) have, on average, the highest number of functional OR genes (854), and the European wolves (n = 4) have the smallest (mean, 801). Mean FORG count in our four European wolf individuals is significantly smaller than that of Asian wolves alone but is not significantly different from that of non-European wolves as a group. Despite having a small repertoire and a sample size of four individuals, the European wolves have, on average, significantly more FORG than domestic dogs if all dog individuals (n=111) are considered (Fig. 1B; Welch t-test, p = 0.007; SI Appendix, Table S1), but this difference becomes non-significant when only dog breed means (n=30) are considered (p=0.17).

Within the domestic dog sample, the number of FORG ranges from 749 to 822 (individual dogs), and 760 to 820 (breed mean). Among breeds, there is a significant variance in gene number (ANOVA, p = 0.004). However, this structure is driven by one breed, the dingo, which has more FORG (820) than any other breed mean (Fig. 1B) and significantly more FORG than three other breeds, the German shepherd, saluki and rottweiler (Tukey HSD pairwise test; p = 0.012, 0.03, 0.01, respectively). When the dingo is removed from the domestic dog sample, there are no significant differences among breeds in mean FORG count (Tukey HSD test, p > 0.11).

We further tested whether concerted artificial selection by breeders has resulted in enhanced FORG repertoires in any breed groupings. First, as regards functional breed groupings, there is no statistically significant difference in FORG repertoire size between non-scent breeds (mean FORG: 778) and scent breeds (mean FORG: 775); (Fig. 1C; Welch t-tests, p=0.37, SI Appendix, Table S1). The lack of difference was present regardless of how we defined scent breeds, as genetically-defined scent breeds alone (3 breeds, 6 individuals; mean FORG: 784), as scent detection breeds (3 breeds, 33 individuals; mean FORG: 773), or as both combined (mean FORG: 775). To test whether years of directed breeding may have favored one sensory specialization over another, we compared scent dogs with sight hounds (Fig. 1C). Scent breeds have, on average, slightly fewer FORG (775) than do sighthounds (786). However, the difference is not significant (Welch t-test; using breed mean, p=0.47; using all individuals within each breed, p=0.08; Fig. 1C). When we extended our test for genetic structure across functional groups to include all ten breed groupings assigned by the American Kennel Club and dog breeders, we again found no significant differences in OR gene repertoire size (Fig. 2A; ANOVA for breed mean, p = 0.9; for individual dogs per breed, p=0.09).

**Figure 2.**
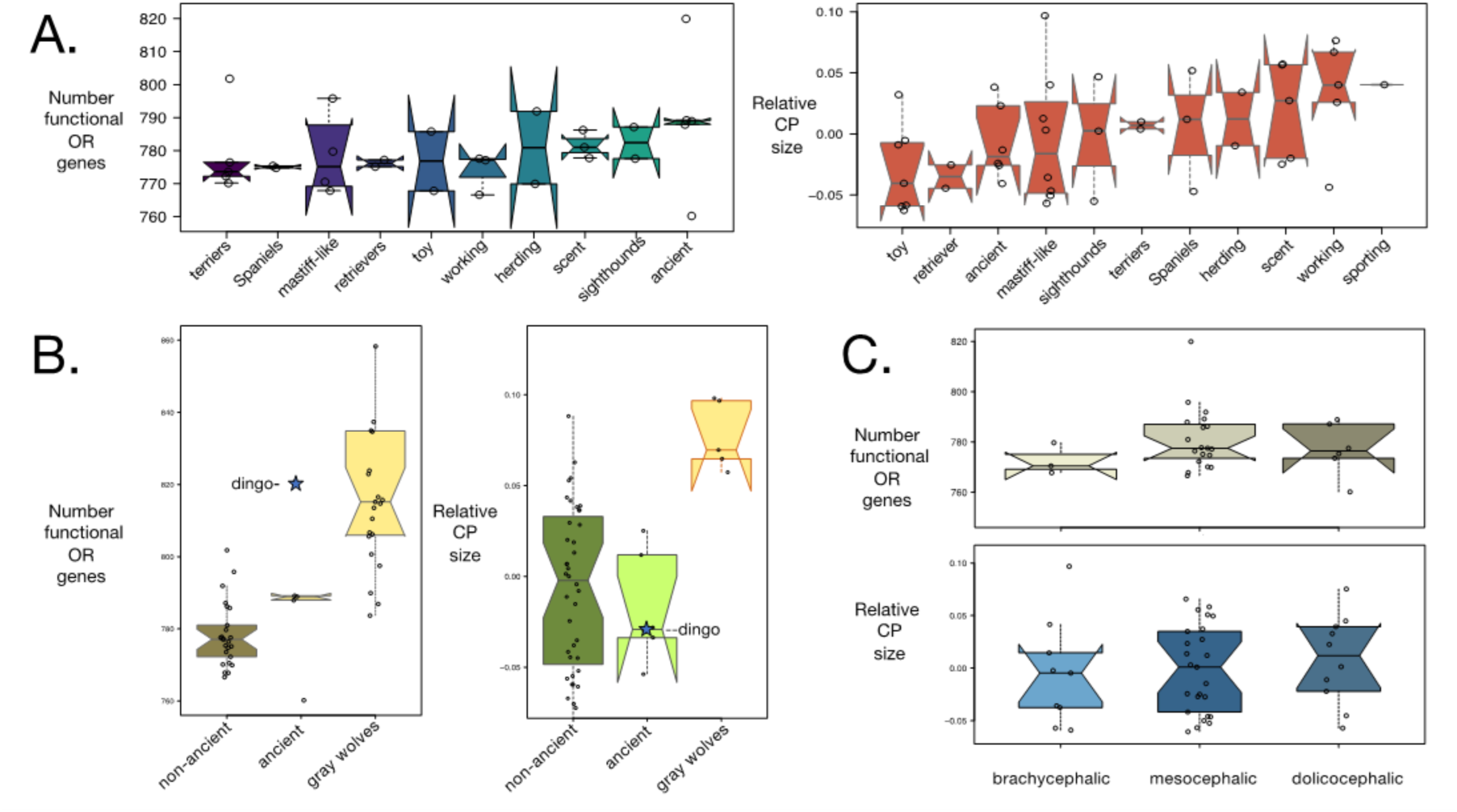
Differential effects of breed groupings on olfactory subgenomes and morphology. A. No significant difference in FORG count (left) and relative CP size (RelCP) (residuals from log-log regression of CP surface area to skull length; see Methods) (right) between the ten dog breed groupings defined by the American Kennel Club and breeders (p=0.9, p=0.6 respectively). B. Ancient dog breeds; (left) mean number of FORG in ancient breeds is significantly different from that of wolves (p=0.012) but not from that of non-ancient breeds (p = 0.31). The dingo FORG count (820) is closer to the mean count in wolves (826) than to that of other ancient dogs (785). Right, the RelCP in the ancient breed grouping is significantly different from that of the wolf but not from that of non-ancient breeds. C. Snout length. No difference in FORG count (upper) and RelCP size (lower) between brachy-, meso- and dolichocephalic dog breeds (p=0.62, p=0.38, respectively). Box plots: midline is median, whiskers are 5%-95% percentile.

Ancient breeds (dingo, basenji, Siberian husky, Afghan, saluki; n = 13 individuals) distinguish themselves from wild canids with a smaller average number of functional OR genes (785) than wolves alone (826) and wolves and coyotes combined (835) (Welch t-test; p=0.014, p=0.012 respectively; Fig. 2B, left). Within domestic dogs, ancient breeds have on average slightly more FORG (785) than do modern breeds (776), however the difference between these two domestic dog groupings is not significant, regardless of whether we test using breed means alone (n=30; Welch t-test, p=0.31) or include all individual dogs (n=111; p=0.12; SI Appendix, Table S1). The dingo is the exception here, with a FORG repertoire (820) closer to the mean FORG count for wolves (826) than to that of other ancient dogs (785) or all other dogs (777) (Fig. 2B).

Finally, as regards morphologically-based breed categories, specifically snout length groupings based on cephalic index (Evans and de Lahunta 2012; Stone et al. 2016), we found no difference in FORG number between brachy-, meso-, and dolichocephalic breeds (Fig. 2C; SI Appendix, Table S2; ANOVA, breed mean, p = 0.62).

### Morphology

To test our hypothesis that differences in olfactory skull morphology across wild and domestic canids would parallel those in olfactory subgenomes, we reexamined and expanded upon cribriform plate data documented in (Bird et al. 2021). With few exceptions, the morphological results parallel the genomic results. In particular, the relative CP size (i.e., size-adjusted CP: residuals from log-log regression of CP surface area vs. skull length) of the domestic dog is, on average, significantly smaller than that of the wolf and coyote combined as well as the wolf alone (N = 53, 51 respectively; ANCOVA; p < 0.0001). Within the wild canids alone, there is no significant difference between coyote and wolf’ relative CP size (ANCOVA; p = 0.25, Fig. 1B, inset left).

Within the domestic dog sample, there are no significant group differences in olfactory anatomy, specifically, relative CP size (RelCP). For example, RelCP size does not differ among domestic dog breeds (Tukey HSD pairwise test; all p-values >0.074) (Fig. 2A, right, Table S2). Moreover, similar to genomic results, there is no difference in RelCP between non-scent breeds and either genetically-grouped scent hounds alone (n=6) or all scent breeds, inclusive of scent detection breeds (n=10) (ANCOVA; p = 0.12, 0.19 respectively Fig. 1D). Similarly, RelCP in scent breeds does not differ from that of sight hounds (ANCOVA; p = 0.25).

Among historical breed groupings, morphological and genomic results differ only slightly. Ancient breeds distinguish themselves from the wild canids by having significantly smaller RelCP than wolves alone as well as wolves and coyotes combined (ANCOVA; p < 0.001, Fig. 2B, Table S1). Moreover, RelCP in ancient breeds (n =6; dingo, basenji, Siberian husky, chow chow, saluki, shar-pei) is not statistically different from that of modern breeds (43). A notable difference between our genomic and anatomical results is that whereas the relative CP size of the dingo is near average for dog breeds, its OR gene repertoire size far exceeds that of all other breeds (Fig. 1B, 2B, SI Appendix, Dataset S1).

Finally, when testing for effects of relative snout length on CP size using direct cephalic index measurements from our skulls, we witnessed no differences in RelCP between brachy-, meso-, and dolicocephalic breeds (ANOVA, breed mean: p = (0.38), Fig. 2C, SI Appendix, Table S2).

### Evolutionary relationship between OR genes and Olfactory morphology

To test whether a previously established correlation between FORG and CP morphology across mammals (Bird et al. 2018) persists within the more recent evolutionary history of dog domestication, we regressed the number of functional OR genes (log10) against RelCP within the wolves, coyotes and all domestic dog breeds for which we have both morphological and genomic data. On the recent time scale of dog breeds alone (N = 20), there is no relationship between functional OR gene number and RelCP (Fig. 3A, SI Appendix, Table S1; r2 = 0.03, p = 0.47). Widening the evolutionary scale to include wolves, then wolves and coyotes, the correlation is reestablished (Fig. 3A, S3, r2 = 0.28, p = 0.006; r2 = 0.37, p < 0.001, respectively). When the wolf and sampled dog breeds are examined in the context of 26 highly-divergent mammal species, the variance among the canids is well within the overall variance, and there remains a strong correlation between OR genes and olfactory morphology (Fig. 3B; r2 = 0.69, p < 0.001).

**Figure 3.**
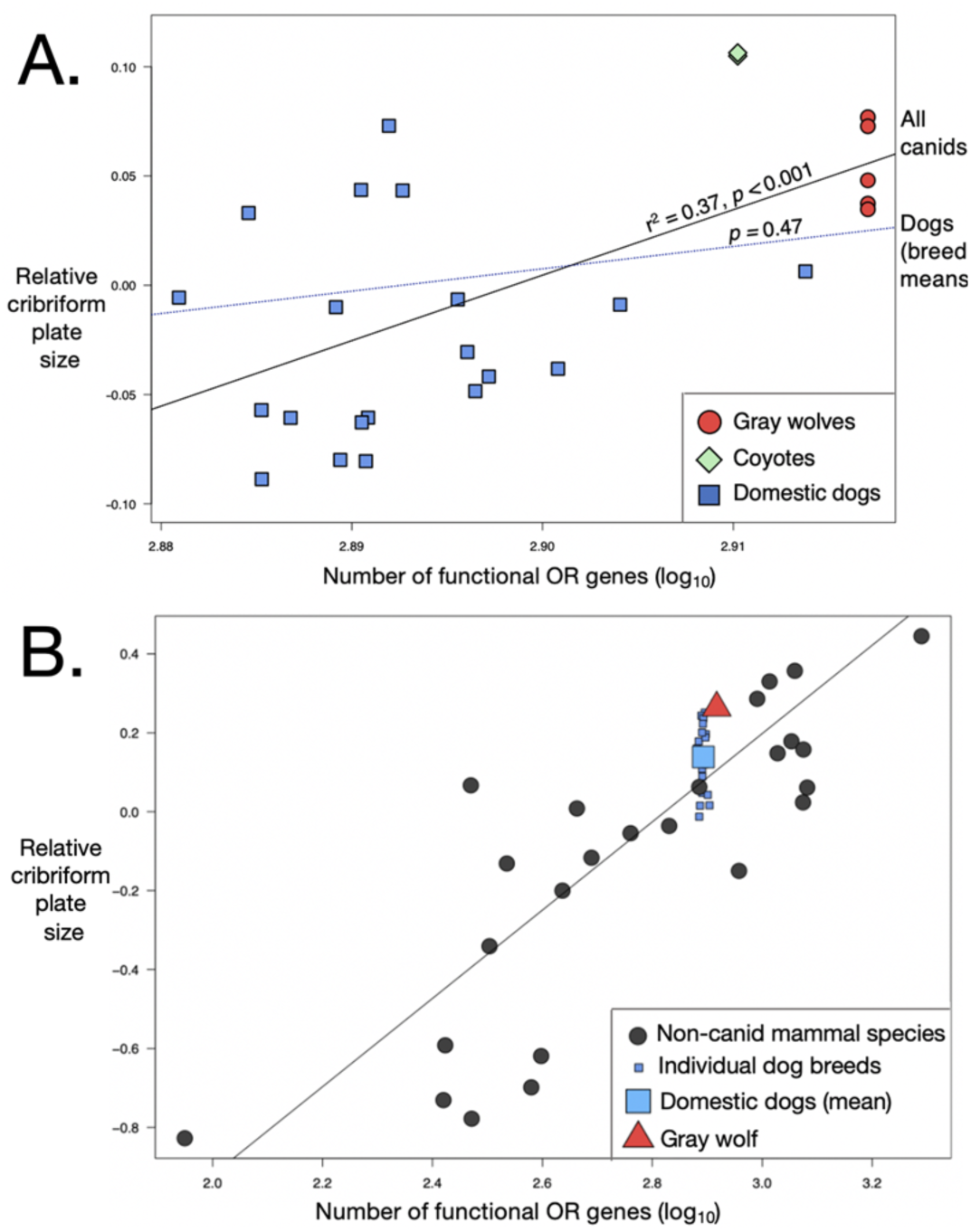
Relationship between relative CP (RelCP) size and number of functional OR genes (FORG) (log10) as a function of evolutionary divergence. A. No significant relationship within the domestic dog breeds (blue) alone. When wolves (red) and coyotes (green) are added, a significant correlation emerges (r2 = 0.37, p < 0.001). B. Addition of RelCP size and FORG data from dogs (breed means, n = 20, small blue squares; species mean from 39 individuals, large blue squares) and gray wolf (species mean from 5 individuals, red triangle) to 26 highly divergent mammal species [black circles, non-canid species means; plot modified from (Bird et al. 2018); Fig. 2b) reveals a strong correlation between RelCP size and FORG repertoires (r2 = 0.69; p < 0.0001). Non-canid mammal species are labeled in Fig. S3.

### Analysis of OR genetic variation

To further investigate whether OR subgenome-wide single nucleotide polymorphisms (SNPs) reveal genetic structure that reflects canid populations, breeds, and functional breed categories, we performed a principal component analysis of 4357 OR SNPs. Domestic dogs, wolves, and coyotes cluster as three discrete groups. The dingo also separates from wolves and dogs on PC1 and PC2 (variance explained 22% and 9%) (Fig. 4A). Principal component analysis, including only domestic dogs and omitting the dingo (to enable better resolution of the dogs), shows two distinct clustering patterns (variance 8% - 7%). First, the ancient breeds tend to be separate from the modern breeds on PC1. Second, the German shepherds form a distinct cluster on PC2 (Fig. 4B). However, there is no noticeable structure between functional groups, such as scent and non-scent breeds or AKC breed grouping.

**Figure 4.**
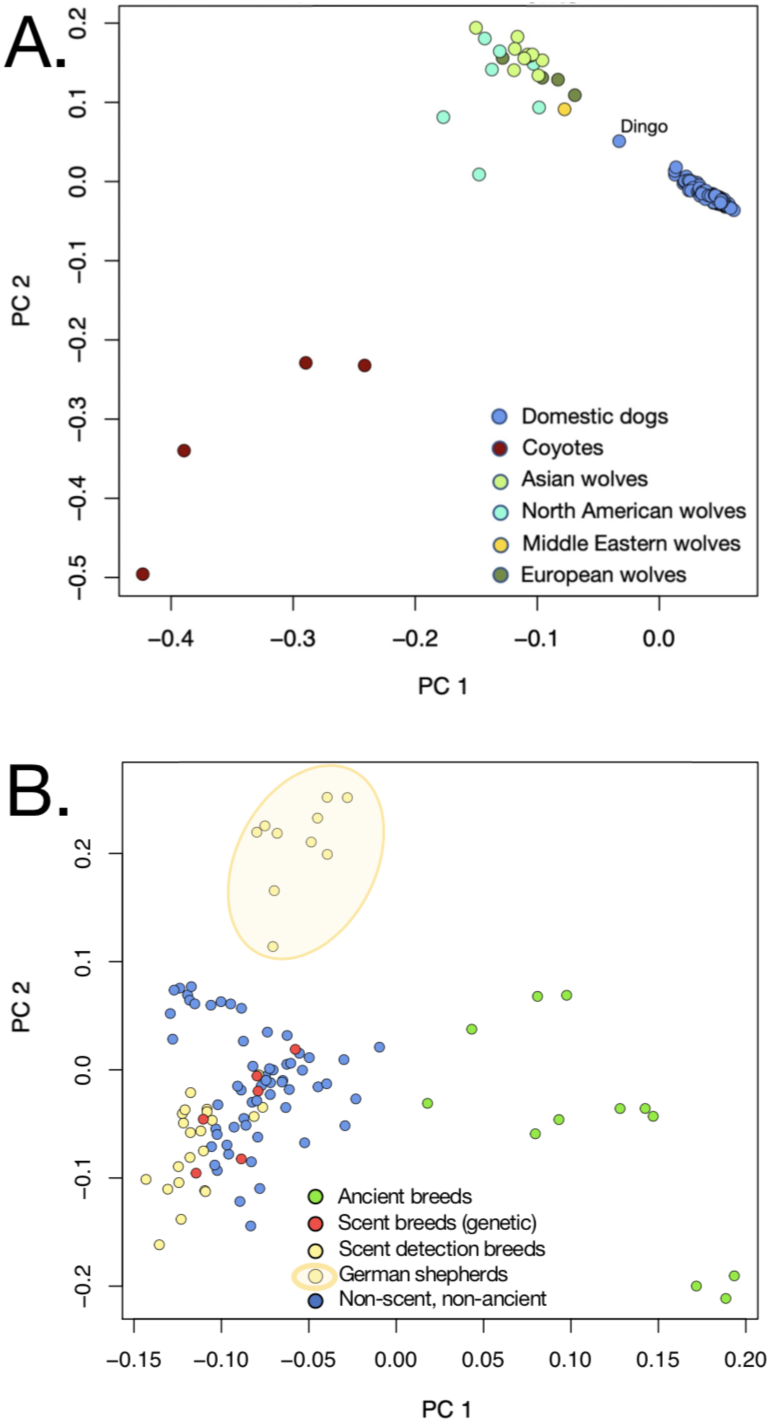
Principal component analysis using the 4357 olfactory receptor (OR) SNPs with each dot representing an individual animal. A. PCA of 29 wolves, four coyotes, and 111 individual dogs shows a clear division between the domestic dog breeds and the wild canids apart from the dingo, which stands separate from both the wolves and dogs on PC1 and PC2. B. PCA of 111 individual dogs shows a separation between the dogs belonging to ancient breeds (green) and those belonging to modern dog breeds on PC1. Modern dogs cluster together regardless of the functional breed grouping on PC1, but German shepherds form a distinct cluster on PC2.

### Gene expression

The RIN scores for the 27 extracts of olfactory epithelium tissues ranged from 5.9 to 9.1 (SI Appendix Dataset S3). One sample was removed as an outlier (S15, code 241132). Hierarchical clustering analyses implemented in WGCNA (Langfelder and Horvath 2008) showed that the biological replicates of RNAseq data from olfactory epithelium clustered well together, and consequently their respective counts were summed for the downstream analyses. After filtering for low counts on the remaining 22 samples (16893 genes) and normalizing the counts, we used Orth in g:Profiler (Reimand et al. 2016) to retrieve the mouse gene symbols. A gene expression matrix for 15138 orthologous genes was then submitted into SaVant (Lopez et al. 2017) (SI Appendix Dataset S4, Fig S1). We removed five samples based on their expression profiles (SI text). After controlling for batch effect, no significant correlation was found with scenting abilities in domestic dogs (SI Appendix, SI text, Dataset S3, Fig. S2).

## Discussion

### Loss of genetic and morphological olfactory capacity in dogs through domestication

Mammals possess species-specific repertoires of olfactory receptor (OR) genes. The number and diversity of OR genes within a repertoire vary markedly across species due to gene gains and losses over evolutionary time associated with distinct olfactory niches and ecological pressures (Niimura and Nei 2007; Hayden et al. 2010; Hughes et al. 2018). Our comparisons of the olfactory receptor subgenomes in 30 breeds of domestic dogs to that of the gray wolf and the closest outgroup, the coyote, determined that domestication has resulted in a significant loss of functional OR genes in dogs relative to wild canines (Fig 1B, SI Appendix, Table S1, Dataset S1). The loss of functional OR genes (FORG) parallels a shift in olfactory skull morphology, specifically to smaller relative cribriform plate size (RelCP) (Fig 1B, SI Appendix, Table S1). Previous studies have established both OR gene repertoire and cribriform plate size as informative molecular and morphological metrics of relative olfactory function, respectively. OR repertoire size is linked to ecological niche (Gilad et al. 2004; Niimura and Nei 2006; Hayden et al. 2010; Niimura 2012; Hayden et al. 2014; Khan et al. 2015; Niimura et al. 2018), ability to discriminate between structurally similar odorants (Laska and Shepherd 2007; Rizvanovic et al. 2013), and the scope of detectable odorants (Malnic et al. 1999; Saito et al. 2009). Likewise, the cribriform plate (CP), a perforated nasal bone that carries a quantifiable imprint of all olfactory nerves on their path from OR cells in the nasal epithelium to the olfactory bulb (Negus 1958; Bird et al. 2014), varies in size across mammals and is linked to habitat and behavioral ecology (Bird et al. 2020). Because RelCP size and FORG repertoire size are strongly correlated across mammalian species (Bird et al. 2018), we investigated this relationship before and after domestication, and in light of artificial selection.

The decline in olfactory metrics in dogs relative to gray wolves suggests that selective pressure for olfactory function has been relaxed in dogs. Although the earliest history of domestication is unclear, dogs likely became increasingly reliant on food sourced from humans as commensal animals, working aides, or pets. In support of this, human and dog diets exhibit parallel shifts in dietary isotopic values over the last 10,000 years (Cannon et al. 1999; Guiry 2012; Sykes et al. 2020). Moreover, coincidental with the expansion of agrarian societies, dogs experienced gene duplications of AMY2B (alpha-amylase), an adaptation to improved starch metabolism (Axelsson et al. 2013). Relative to wild canines, dogs generally do not locate and track prey over large home ranges, and losses in olfactory capacity may reflect relaxed selective constraints on maintaining an extensive gene repertoire (David Mech 1966; Gittleman 1991). Alternatively, a reduced olfactory function might be a passive result of drift-related olfactory gene diversity loss due to at least two bottleneck events, first during domestication and later during breed formation (Wayne and Ostrander 2007; Cruz et al. 2008; Freedman et al. 2014). We detected no significant differences between coyote and wolf FORG repertoires but did recover significant variance among wolf populations. Among the wolves in our sample, those from Europe have, on average, the smallest repertoire, while Asian wolves have the largest. The FORG disparity among wolves may have its roots in ancient population structure. Following the divergence of New and Old-World wolves (ca. 11-12 kya), European wolf lineages, particularly Southwestern European, experienced a marked drop in effective population size relative to other wolf populations, likely due to a severe demographic bottleneck (Lucchini et al. 2004; Sastre et al. 2011; Freedman et al. 2014; Pilot et al. 2014; Fan et al. 2016; Hulva et al. 2018). Long-term isolation and bottleneck events in the European wolves could have led to the contraction of superfamily genes, suggesting that below a certain population size, balancing selection may not be strong enough to maintain genetic diversity (Quignon et al. 2005; Ploshnitsa et al. 2012). However, genetic variability of immunity-related genes seems to have been preserved in European wolves despite these bottlenecks (Arbanasić et al. 2013; Galaverni et al. 2013; Niskanen et al. 2014). Notably, there is a sizable variance within our sample of four European wolves. The three individuals from the Iberian Peninsula and Italy, a population known for long-term isolation, have a particularly low number of functional OR genes (mean = 789). By contrast, the single individual from Croatia has 835 functional OR genes, commensurate with the average FORG count for all the wolves in our sample (826). While all wolf lineages experienced a decline in effective population size, Asian wolves, except Tibetan populations, do not appear to have undergone bottlenecks as severe as those of European wolves. Chinese wolves seem to have experienced both marked population growth and decline during the Late Pleistocene (Fan et al. 2016), which may help explain the high variance in FORG number among our sample of Asian wolves. However, clarification of why Asian wolves exhibit greater variance and a larger mean count of FORG, would require a targeted study on the evolution of the OR repertoire in different populations of wolves based on a high-quality wolf reference genome, a task beyond the scope of this study. Because our analyses are based on a dog genome reference assembly (from a boxer), we were unable to detect copy number variants of wolf-specific gene families.

### Conditional relationship between OR gene number and olfactory morphology

Previous work established a strong linear correlation between the number of functional OR genes (FORG) and relative cribriform plate size (RelCP) among 26 species representing all mammalian superorders, ranging in body mass from 0.1 kg to over 2900 kg, and including the domestic dog (Bird et al. 2018). Here, we asked if this morphologic-genetic relationship is also supported on a more recent evolutionary timescale and within a species, specifically in canine populations that diverged as recently as 15 kya. While some breeds, like the dingo, husky, and basenji diverged earlier in the process of dog domestication, most dog breeds have diversified very recently in the Victorian era beginning about 200-300 years ago with the advent of selective breeding (Parker et al. 2017). We found no significant linear correlation between FORG repertoire and RelCP among domestic dog breeds alone. However, adding wolves and coyotes to the analysis restored the relationship (Fig. 3A). Because of the recent development of most dog breeds and crossbreeding, the correlation between FORG and olfactory morphology may only be apparent when more divergent lineages, such as the wolf and other mammal species, are included. Indeed, when canids are added to the divergent group of mammal species included in the previous study (Bird et al. 2018), the relationship between FORG and RelCP size among wolves and dog breeds is well within the overall variance across mammal species, and the overall linear correlation remains strong (r2 = 0.69, p < 0.001) (Fig. 3B). It is worth noting that across dog breeds alone, variance in RelCP size is larger than variation in FORG number. Because we know that CP shape is informed by extreme skull shape differences in dogs (Jacquemetton et al. 2021), it is conceivable that CP size is also influenced by the profound variation in dog snout size and shape (Schoenebeck and Ostrander 2013) and that directed artificial selection on snout size and skull phenotype has weakened the relationship between CP size and number of OR genes.

### Dog breeds and olfaction: scent hounds in name alone

A central finding in this study was that scent hounds show no expansion in the number of FORG relative to non-scent breeds. Similarly, there is no sign of olfactory enhancement in scent hounds relative to sight hounds. These findings hold whether the scent grouping is made up solely of those breeds defined as a monophyletic clade of scent dogs (beagle, bloodhound, dachshund) (Vonholdt et al. 2010; Parker et al. 2017) or if it includes breeds preferentially used as scent detection dogs (German shepherd, Labrador retriever, golden retriever) (Ensminger 2011; Rocznik et al. 2015). Morphological findings parallel that of gene diversity, in that the relative size of the cribriform plate in scent dogs is no larger than that of non-scent dogs or even sight hounds (Fig. 1C, SI Appendix, Table S1, Dataset S1) (Bird et al. 2021).

A surprising pattern emerged in the principal component analysis of OR gene SNPs among dogs alone. German shepherds, defined here as scent detection dogs, cluster separately from both the non-scent and scent dogs (Fig. 3B). At this point it is difficult to determine whether this distinct clustering is due to an earlier demographic event, possibly a bottleneck, in the history of the German shepherd breed, and/or whether it represents a functional difference. We note here that all German shepherd dogs in our sample underwent unique gene losses in an OR gene cluster on Chromosome 21 (Chr21:26733324-26734268, Chr21:26751570-26752529), however it is beyond the purview of this paper to determine whether this loss is tied to the pattern we see among the OR gene SNPs.

To gain further insight into the specific interaction of genes that might affect olfactory performance, we assessed patterns of OR gene expression between scent and non-scent breeds. Gene regulatory mechanisms allow a range of phenotypes to arise from an otherwise static genome sequence (82). However, we did not find any significant association between gene expression and olfactory function in dog breeds. We have to acknowledge that this conclusion might be limited by the small number of individuals and unique breeds used in the analysis as well as the possibility that not all OR genes were retrieved during the sampling process.

Overall, we found no morphologic or genetic evidence that breeds categorized as scent hounds are superior smellers or were bred specifically for olfactory ability. Our results challenge claims by breeders that olfactory traits have been selected and managed through strict controls over reproduction among scent breeds (Pemberton 2013). Despite the elevated status given to the best-known scent hound, we found that the bloodhound sits squarely in the middle of domestic dog breeds, both in OR gene repertoire and olfactory anatomy (Figs. 1B, SI Appendix, Fig. S6). To illustrate our struggle to find data that support the acclaimed olfactory ability of scent hounds, we constructed a graphic analysis of a cascade of unsupported references in the primary literature that repeat the misconception that the bloodhound and other scent hounds have an unparalleled olfactory anatomy (SI Appendix, Fig. S7).

### The dingo and ancient dog breeds

“Ancient dogs’’ comprise a group that is genetically highly divergent from other dogs and includes breeds that originated from ancient cultures >∼500 years ago and show strong evidence of admixture with wolves after their domestication (Vonholdt et al. 2010; Freedman et al. 2014; Parker et al. 2017). Ancient breeds in our sample include the dingo, basenji, Siberian husky, saluki, and Afghan hound. On average, the ancient breed grouping has a higher number of functional OR genes than modern breeds and a lower number than the wild canids. (Fig. 1B, 2B). However, statistically, the ancient dogs, as a group, align with the modern dogs and differentiate themselves from the wolves and coyotes (SI Appendix, Table S1). The dingo is the exception, with an FORG repertoire closer to that of the wolves than to that of other ancient dogs. Dingos belong to a genetically divergent group of domestic dogs isolated for thousands of years (Freedman et al. 2014; Field et al. 2022). Compared with dogs, dingoes are considered feral and hunt prey independently from humans (Savolainen et al. 2004; Zhang et al. 2020). Because they are less reliant on human food sources than dogs consuming starch-rich diets since the Neolithic, dingos likely did not experience relaxed selection on olfactory specialization. Consistent with these observations, the dingo has retained the wolf-like condition of a single copy of AMY2B in contrast to other dog breeds (Arendt et al. 2016; Field et al. 2022). However, it is noteworthy that the dingo’s high number of functional OR genes is not reflected in a larger cribriform plate (Fig. 2B).

### Breed groups and olfaction

The spectrum of variation in the number of functional OR genes is relatively modest across dog breeds and does not reflect breed grouping. Dog breeds have historically been grouped according to function and genealogical relationships (Wilcox and Walkowicz 1989; American Kennel Club 2007; Judah 2007; Vonholdt et al. 2010; Parker et al. 2017). Breed groupings used today by breeders and the American Kennel Club are comprised of the ancient, spitz, toy, spaniels, scent hounds, working dogs, mastiff-like breeds, small terriers, retrievers, herding, and sight hounds (Vonholdt et al. 2010). Although these common classifications have fairly modest genetic support as revealed by haplotype-sharing and allele-sharing analyses (Vonholdt et al. 2010; Parker et al. 2017), they persist throughout the literature.

Our investigations of functional OR gene count show no significant differences among commonly used breed groupings (Fig 2A). Notably, sight hounds have a slightly higher number of FORG than scent hounds (Fig 1C), contradicting the notion that sight hounds were selected for their visual abilities, whereas scent hounds were selected for their superior noses. Morphological comparisons of cribriform plate size revealed the same lack of significant distinctions across common breed groupings (Fig 2A). Olfactory gene SNP analysis among dog breeds failed as well to reveal differences between common groupings, except for the ancient breeds, which may be driven by the dingo. We expected that there might be patterns of sensory specialization that matched the AKC functional groupings, given the evidence of positively selected genes associated with athletic ability in sport-hunting breeds (Kim et al. 2018), as well as a selected trade-off between limb bone strength and stiffness in the American pit bull terrier and greyhound (Kemp et al. 2005). AKC breed groupings undoubtedly have some basis in the history of directed breeding for function, however this does not appear to apply to olfaction.

### Olfaction and snout length

Skull shape has been a central focus of artificial selection throughout the domestication of dogs, resulting in a continuum of snout lengths encompassing that exhibited in wolf ontogeny (Wayne 1986; Wilcox and Walkowicz 1989; Drake and Klingenberg 2010; Drake 2011; Schoenebeck and Ostrander 2013; Georgevsky et al. 2014). Most domestic dog breeds have shorter faces (palates) than the wolf, the most pronounced of which are found among the brachycephalic dogs (Pekingese, pug, collective bulldogs). On the other end of the continuum are the long-faced, dolichocephalic breeds (saluki, collie, borzoi). Short-snouted breeds are not known for their olfactory performance and are generally avoided by detection dog trainers (Jamieson et al. 2017); however behavioral studies of detection performance in brachycephalic dogs relative to non-brachycephalic dogs show contradictory results (Polgár et al. 2016). Here, we found no significant difference in OR repertoire size among brachycephalic breeds relative to both meso- and dolichocephalic breeds (Fig 2C) suggesting that individual differences in olfactory performance among brachycephalic dogs might reflect structural constraints imposed on nasal anatomy by positive artificial selection for short-snoutedness. Short-faced dogs are prone to respiratory and upper airway syndromes (Lorinson et al. 1997), which affect airflow and may conceivably influence olfactory function. Notably, in our morphological analysis, there was relatively high variance in RelCP across brachycephalic individuals, however there was no difference in the RelCP between brachy-, meso- and dolichocephalic breeds (Fig 2C). Therefore, relative snout length based on the cephalic index we used here, ratio of maximum skull width to skull length, appears to be a poor predictor of olfactory morphology or gene diversity. However, it is conceivable that a relative snout size metric different from cephalic index may better describe how selection for snout size and shape has regulated the expansion of the olfactory skeleton and innervation.

In summary, our results indicate that relative to their closest living relatives, gray wolves and coyotes, domestic dogs have a reduced OR subgenome and olfactory skeleton. Within domestic dogs, ancient breeds do not appear to have retained ancestral or wolf-like olfactory attributes relative to modern breeds. One exception is the dingo, which has a larger number of functional OR genes than any dog breed in our sample. We found no evidence of direct selection for an elevated sense of smell among scent breeds. Contrary to popular belief that scent hounds have superior noses, our results reveal that scent breeds are not distinguished from other dog breeds in either OR gene repertoire, OR gene expression or relative cribriform plate size. Artificial selection for short faces in brachycephalic dogs has not resulted in any significant reduction of the olfactory variables we measured. Overall, there is considerable variability within breeds in the number of functional olfactory genes; however, no breed grouping stands out, suggesting that most or all dogs can perform olfactory based functions. The apparent ability of some breeds to perform scent detection tasks better than others likely reflects aspects of behavior, such as motivation and trainability, rather than olfactory gene repertoire and anatomy.

## Materials and Methods

### Morphometry Sampling

We sampled 103 skulls from 45 identified dog breeds, one unknown dog breed, and two species of wild canid, gray wolf (*Canis lupus*) and coyote (*Canis latrans*) (SI Appendix, Table S1). All specimens were sourced from museum and university collections listed in SI Appendix, Table S1. Sampled wild canid species include only wild-caught adult specimens. Species and breed body masses, as estimated from the literature (Nowak 1991; Crowley and Adelman), ranged from approximately 2.25 to 68 kg.

### Genomic Sampling

OR gene copy number variation was estimated from 111 domestic dog genomes belonging to 30 different breeds. To make domestication-based inferences, we compared dog repertoires to those estimated from the following wild *Canis* genomes: 27 gray wolves (19 old world wolves and 8 New World wolves), and 4 coyotes (SI Appendix, Dataset S1).

### Gene Expression Sampling

Dogs that were admitted to the Texas A&M University Veterinary Medical Teaching Hospital and euthanized by owner request were included in this study. Animals with a history and physical assessment indicative of nasal or upper respiratory disease or infectious disease were excluded. All samples were acquired within 2 hours of euthanasia. The temporal horns of the frontal sinuses were identified by surface palpation and a region on midline 1-2 cm rostral to the temporal horns was selected for trephine. A 1-2 cm elliptical skin incision was made on midline and soft tissues dorsal to the nasal and frontal bones were dissected (SI Appendix Fig. S8). A 4 mm diameter sterile trephine was used to remove a round section of bone, overlying the cribriform plate. Sterile forceps were used to grasp multiple pieces of mucosa immediately underlying the punch as well as 5 mm rostral, caudal, and abaxial to the trephine.

### Breed identification and sample size

Breed type was assigned by original dog owners or museum collectors. Where possible we sampled two or more individuals per breed, preferably from each sex. We recognize that sample size per breed is relatively low, however given the large number of breeds and species in our study, a deeper sampling was prohibitive (limited number of specimens and quality dog genomes available).

### Breed groupings

We used four criteria to classify domestic dogs into breed groupings. First, we grouped the breeds into (a) scent breeds (Vonholdt et al. 2010; Ensminger 2011; Rocznik et al. 2015; Parker et al. 2017) and (b) non-scent breeds. Within this classification, the scent breeds are divided into two sub-groups: genetically defined scent breeds, that is, those that have been defined as a monophyletic clade in molecular studies (beagle, bloodhound and dachshund (see (Vonholdt et al. 2010; Parker et al. 2017) and detection dogs, that is, breeds outside that clade that are commonly chosen for scent detection work (German shepherd, golden retriever, and Labrador (Ensminger 2011; Rocznik et al. 2015). This latter classification was used for the gene expression analyses as well. Second, in classifying dogs as ancient and modern breeds, we used criteria used by (Vonholdt et al. 2010; Freedman et al. 2014; Parker et al. 2017) to identify the ancient breeds in our sample (dingo, basenji, Siberian husky, saluki and Afghan hound). Third, we classified dogs into ten functional breed groupings traditionally used by breeders and the American Kennel Club (Wilcox and Walkowicz 1989; American Kennel Club 2007; Vonholdt et al. 2010). Finally, we used relative snout length, specifically the cephalic index (CI; ratio of maximum skull width to skull length x 100) (Roberts et al. 2010; Evans and de Lahunta 2012) to group most dogs into brachy-, meso- and dolichocephalic breeds. Because we had no access to the skulls of the individual dogs represented in our genome data, we used mean breed CI values already established in an extensive study by (Stone et al. 2016).

### Data collection Morphology

All skulls were scanned on high-resolution industrial computed tomography (CT) scanners (Phoenix v|tome|x S; North Star Imaging ACTIS; XRadia MicroXCT; Nikon Metrology XT H 225 ST). The targeted region of interest was constrained to the CP and the area directly surrounding it in order to increase scan resolution. Scan voxel size ranged from 0.04 mm to 0.085 mm. All scan data are available through MorphoSource (https://www.morphosource.org/) or Digimorph (http://www.digimorph.org). To visualize and quantify CP morphology, we imported CT scan data into the 3D imaging software Mimics (v. 20.0-21.0, Materialise Leuven, Belgium), segmented the CP into masks that delineate bone and non-bone, and finally reconstructed 3D volumetric models (Fig 1A, SI Appendix, Movies S1,2). CP surface area is defined here to include only the area of bone perforated by foramina that surround olfactory nerves, a proxy for the amount of olfactory innervation found in an animal’s snout. Previous work established a strong linear relationship between the cumulative surface area of the CP foramina and the surface area of the perforated portion of the CP (Bird et al. 2014). This excludes the lateral flanks of the CP perforated by the ethmoid foramen, a distinctly large passageway for the nasociliary branch of the trigeminal nerve that has no olfactory function. We quantified CP surface area first by rendering the perforated area into a continuous surface area in the imaging program 3-matic (v. 11.0-13.0, Materialise) with a wrapping function that fills all foramina in the CP model and then second, by digitally incising the CP surface along the perimeter of the perforated region (SI Appendix, Fig. S9). We digitally calculated the surface area in 3-matic.

### RNA extraction

Total RNA was extracted from 24 epithelium tissues using the Invitrogen TRIzol® Plus RNA Purification Kit. Four samples were extracted in two separate batches (SI Appendix, Dataset S3). The integrity of 28 RNA extracts was then quantified using the Agilent bioanalyzer (Agilent Technologies, USA). One sample (code 225627) was removed from the library preparation due to low RNA integrity number (RIN) score (2.4). The RIN scores for the remaining extracts ranged from 5.9 to 9.1 (SI Appendix, Dataset S3). cDNA libraries were constructed using the KAPA mRNA HyperPrep Kit with dual indices (Kapa Biosystems, Ltd). Individual libraries were then pooled in equimolar ratios and sequenced on two lanes of an Illumina Hiseq4000 (150bp paired-end). Sequencing was performed at Fulgent genetic (https://www.fulgentgenetics.com).

### Data analyses Morphology

Because CP area increases with body size, we calculated a metric of size-adjusted relative cribriform plate size (RelCP). This size-corrected metric was estimated following (Bird et al. 2018) using residuals from an ordinary least squares regression of log10 values of absolute CP surface area against a body size proxy for all breeds and the two wild canid species. As a body size proxy, we used the distance between the occipital condyles and the anterior extent of the orbit (OOL, occiput to orbit length), a cranial metric shown to correlate well with body mass in carnivorans (r2 = 0.9)(99) (SI Appendix, Fig S10). In our overall analyses we chose OOL over total skull length or body mass as a size proxy for two reasons. First, OOL excludes snout length and avoids the confounding effects of large variation in snout length (i.e. brachycephaly and dolichocephaly) present in our sample of dog breeds (Schoenebeck et al. 2012). Second, weight was not available in collectors’ notes for most specimens, and weights reported by the American Kennel Club are based on breed averages and display large ranges (Crowley and Adelman). A log-log generalized least squares regression of mean absolute CP surface area against OOL was used to derive RelCP from resulting residuals. In a single case, when analyzing the relationship between OR genes and CP plate morphology within a wider context of highly diversified mammals with variable skull morphologies (SI Appendix, Fig. S6), we used body mass as a size correction for CP size in order to match the original study of (Bird et al. 2018). Phylogenetic comparative methods were not used here to account for the effects of phylogeny on CP morphology, as existing cladograms for wolves and domestic dog breeds are not time-calibrated due to extensive admixture between breed lineages (Parker et al. 2017). To test for significant differences in RelCP between wild canids and domestic dogs and between dog breed groupings, we performed pairwise t-tests and one-way analysis of variance (ANOVA). Additionally, while testing for differences in RelCP between groupings in various subsets of the data, we performed an analysis of covariance (ANCOVA), as it is robust to violations of normality. We carried out all analyses in R (v. 3.5.3) (R Core Team, 2014).

### Genome mapping and SNP calling

We applied Trim galore (Trim Galore, http://www.bioinformatics.babraham.ac.uk/projects/trim_galore/) to filter paired-end Illumina reads. The trimmed paired-end reads were mapped to the domestic dog genome assembly (Version CanFam3.1) using BWA-mem (Li and Durbin 2010) with default settings. SAMtools (Li et al. 2009) was used to remove PCR-induced duplicates. The standard Genome Analysis Toolkit (GATK) (Van der Auwera et al. 2013) pipeline was used for base quality recalibration and indel realignment.

### OR gene annotation of dog genome assembly

To improve the accuracy of OR gene annotation of the dog genome assembly CanFam3.1, we applied a modified Perl script pipeline (https://github.com/GanglabSnnu/OR_identify) to identify all intact (functional) and pseudogene OR genes (Montague et al. 2014). We define functional OR genes as those meeting the following criteria: (i) no premature stop codon, (ii) no frameshift mutations, (iii) no in-frame deletions within a single transmembrane region nor deletions of conserved amino acid sites (Niimura 2013), (iv) no truncated genes with fewer than 250 amino acids or lacking any of the seven transmembrane domains (Hayden et al. 2010). The updated OR gene annotation was used to estimate OR gene copy number variation (CNV) on all 142 canine genomes.

### SNP and Copy number variant (CNV) analyses

The software snpEff (Cingolani et al. 2012) was applied to annotate different categories of SNP and INDEL variants (e.g., premature stop codon, frameshift, synonymous substitution and non-synonymous substitution). We use a consensus from two structural variant callers, CNVnator (Abyzov et al. 2011), and Delly2 (Rausch et al. 2012) to estimate the CNV of all intact and pseudogene of OR among all analyzed dogs and wolves’ individuals. We performed principal component analysis (PCA) with the R package ggfortify to visualize the relationships of OR gene copy number and substitution variants among all canid species.

### OR Gene expression analyses

Raw sequences were processed using Trim Galore 0.3.1 (Krueger) to remove Illumina adapters and sequences that did not meet the following quality thresholds: Q > 20, length > 25). The alignment of the trimmed reads was performed on STAR 2.5.3 (Dobin et al. 2013) using the dog genome (*Canis lupus* familiaris: Ensembl release 95_31). We used HtSeq for read counts on a custom GTF file including all intact olfactory genes. We first checked for biological replicates and outliers. We filtered reads with low counts in the 27 samples and remaining genes were normalized using TMM (trimmed mean of M-values) in the edgeR package (Robinson and Oshlack 2010) in R. Reads were then converted into log2 counts per million (logCPM) with voom in LIMMA (Law et al. 2014; Ritchie et al. 2015). We performed principal components analyses to identify technical factors from the dataset (Blighe and Lun 2020). We removed the batch effects using the removebatch command in LIMMA for visualization purposes. We explored the data to check for outliers and clustering of biological replicates using hierarchical clustering of the gene expression adjacency matrix with the R package WGCNA (Langfelder and Horvath 2008).

After controlling for biological replicates and outliers, we checked for tissue-type heterogeneity using a web-based tool named SaVant (http://newpathways.mcdb.ucla.edu/savant-dev/) (Lopez et al. 2017). SaVant accepts a matrix of gene expression from RNAseq or microarray and allows a comparison between our own expression data with a repository of more than 10895 signature profiles. Exploration of expression profiles for the remaining samples was investigated for known olfaction signatures such as “Olfactory bulb”, “Kegg olfactory transduction” (389 genes), and “Reactome olfactory signaling pathway” (328 genes). We performed an orthology search of the dog genes with the mouse symbol genes using:Orth in g: Profiler (Reimand et al. 2007; Reimand et al. 2016). We suggest that RNA samples with a weak to nonexistent OR signature were likely due to sampling bias (lacking olfactory epithelium) and were removed from the analyses.

### Differential expression and gene enrichment analyses

RNAseq reads were filtered and processed as explained above. LIMMA (Law et al. 2016) and Deseq2 (Love et al. 2014) were used for our differential gene expressions analyses. Genes falling below FDR<0.05 in both methods were kept for Gene Ontology analyses in g:Profiler (Reimand et al. 2016). To ensure that the distribution of the false discovery rate satisfied the expectation for FDR under a null model, we ran 100 permutations of the original model where the significant variable was assigned randomly and compared the distribution of p-values under the null to the distribution of p-values under the true model.

## Acknowledgments

This work was supported by National Science Foundation grants IOS-1457106 to R.K.W. and B.V.V. and IOS-1456506 to W.J.M. and J.L. A.M. and D.B. were supported by the NSF grant (IOS-1457106). A.M. and M.M. were supported by the QCBio Collaboratory Postdoctoral Fellowship (UCLA). A.M. used computational and storage services associated with the Hoffman2 Shared Cluster provided by UCLA Institute for Digital Research and Education’s Research Technology Group. We thank curators and collection managers: M. Flannery of California Academy of Sciences, C. Conroy of Museum of Vertebrate Zoology, UC Berkeley, K. Molina of the Donald R. Dickey Collection, J. Dines of the Museum of Natural History Los Angeles County, K. Zyskowski of Yale Peabody Museum for providing skulls; M. Colbert, R. Ketchum, J. Maisano of University of Texas HRCT Digital Morphology group, T. Jashaashvili, T. Skorka of Keck MIC, M. Faillace, J. Urbanski of General Electric Inspection Technologies, and T. Stecko, T. Ryan of Pennsylvania State University for producing high-quality CT scans. We thank Elizabeth Scanlan (TAMU) for domestic dog tissue sampling and preservation.

## Conflict of Interest

The authors declare no conflicts of interest

## Data availability

The raw sequencing data, normalized counts, regressed normalized counts, and all associated metadata have been deposited in NCBI’s Gene Expression Omnibus and are accessible through the GEO Series accession numbers (xxxx). Data from CT scanning will be available on Morpho Source (https://www.morphosource.org/).

## Notes

### Competing Interest Statement

The authors have declared no competing interest.

