## Supplementary material for "Genetic and anatomical determinants of olfaction in dogs and wild canids"

### Figures

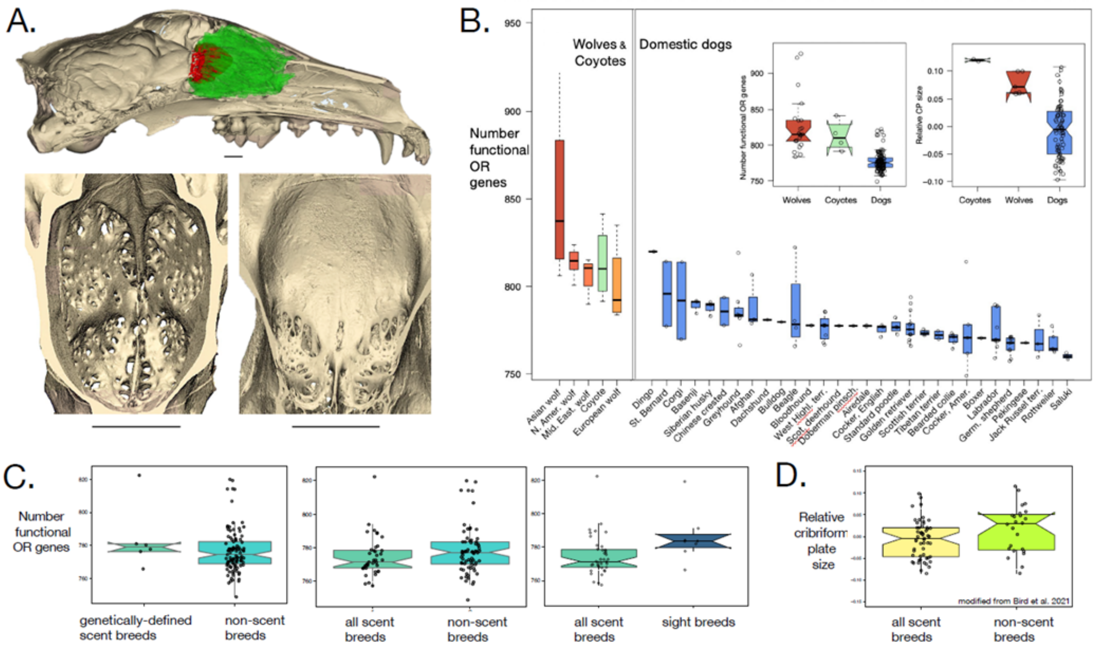


**Figure 1.** Comparative olfactory morphology and genomics in wild canids and dog breeds. A. Morphological metric, cribriform plate (CP) in skull matrix; Top, Borzoi half skull, sagittal view; red, CP; green, olfactory turbinal bones. Bottom, CP, posterior view, showing the relative presence of foramina (holes) for olfactory nerve passage in the gray wolf (left) and Pekingese (right). Scale bars, 10mm. B. Number of functional olfactory receptor genes (FORG) in wolves (orange), coyotes (green), and domestic dog breeds (blue) in descending order. Left inset, Domestic dogs have a smaller FORG repertoire than gray wolves and combined wolves and coyotes ($p<0.001$). Right inset, relative CP (RelCP) size in dogs is, on average, smaller than in wild canids ($p<0.001$). C. Left, FORG count in genetically-defined scent breeds is not significantly different from non-scent breeds ($p=0.41$). Middle, Mean FORG count for all scent breeds is not significantly different from that of non-scent breeds ($p=0.16$) and (right) sight breeds ($p=0.08$). D, RelCP size is no different between scent and non-scent breeds ($n=46$, $p=0.12$). Box plots: midline is median, whiskers are 5%-95% percentile.

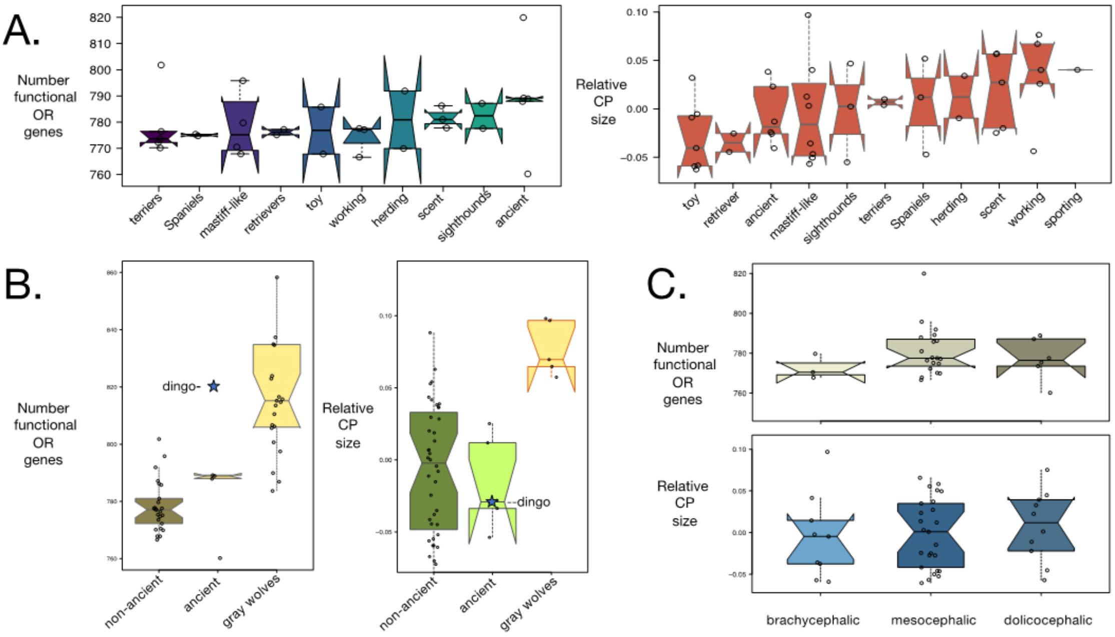


**Figure 2.** Differential effects of breed groupings on olfactory subgenomes and morphology. A. No significant difference in FORG count (left) and relative CP size (RelCP) (residuals from log-log regression of CP surface area to skull length; see Methods) (right) between the ten dog breed groupings defined by the American Kennel Club and breeders (p=0.9, p=0.6 respectively). B. Ancient dog breeds; (left) mean number of FORG in ancient breeds is significantly different from that of wolves (p=0.012) but not from that of non-ancient breeds (p = 0.31). The dingo FORG count (820) is closer to the mean count in wolves (826) than to that of other ancient dogs (785). Right, the RelCP in the ancient breed grouping is significantly different from that of the wolf but not from that of non-ancient breeds. C. Snout length. No difference in FORG count (upper) and RelCP size (lower) between brachy-, meso- and dolichocephalic dog breeds (p=0.62, p=0.38, respectively). Box plots: midline is median, whiskers are 5%-95% percentile.

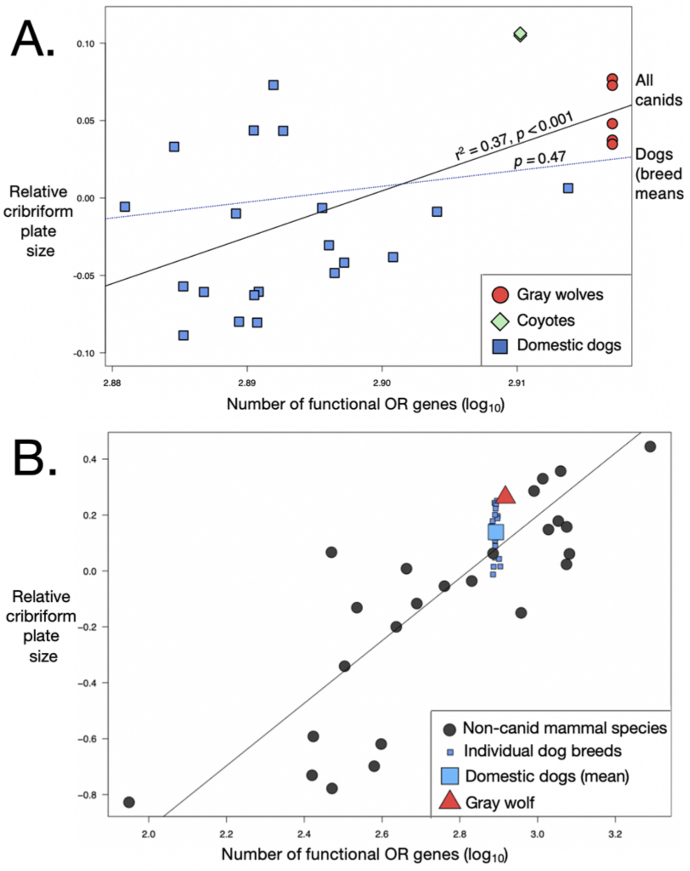


**Figure 3.** Relationship between relative CP (RelCP) size and number of functional OR genes (FORG) (log10) as a function of evolutionary divergence. A. No significant relationship within the domestic dog breeds (blue) alone. When wolves (red) and coyotes (green) are added, a significant correlation emerges (r2 = 0.37, p < 0.001). B. Addition of RelCP size and FORG data from dogs (breed means, n = 20, small blue squares; species mean from 39 individuals, large blue squares ) and gray wolf (species mean from 5 individuals, red triangle) to 26 highly divergent mammal species [black circles, non-canid species means; plot modified from (Bird et al. 2018); Fig. 2b) reveals a strong correlation between RelCP size and FORG repertoires (r2 = 0.69; p < 0.0001). Non-canid mammal species are labeled in Fig. S3.

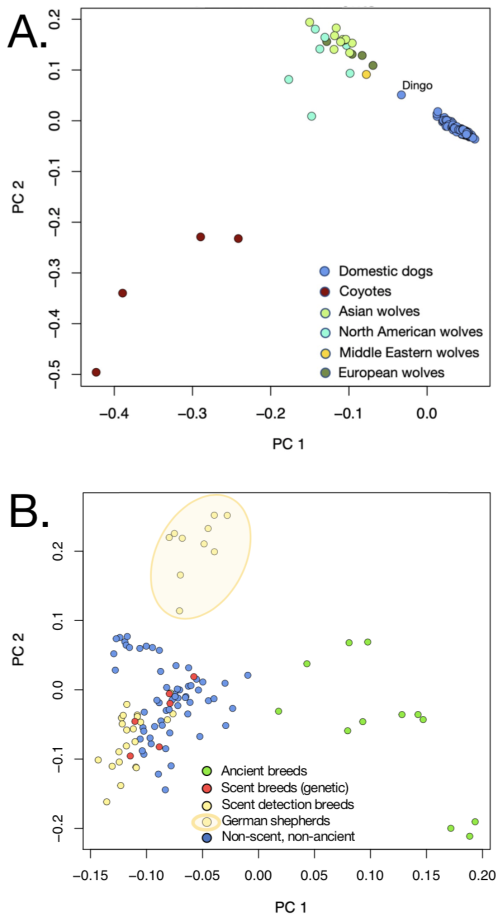


**Figure 4.** Principal component analysis using the 4357 olfactory receptor (OR) SNPs with each dot representing an individual animal. A. PCA of 29 wolves, four coyotes, and 111 individual dogs shows a clear division between the domestic dog breeds and the wild canids apart from the dingo, which stands separate from both the wolves and dogs on PC1 and PC2. B. PCA of 111 individual dogs shows a separation between the dogs belonging to ancient breeds (green) and those belonging to modern dog breeds on PC1. Modern dogs cluster together regardless of the functional breed grouping on PC1, but German shepherds form a distinct cluster on PC2.
